# Phage lysis protein Lys^M^ acts as a wedge to block MurJ conformational changes

**DOI:** 10.1101/2025.04.26.650401

**Authors:** Hidetaka Kohga, Napathip Lertpreedakorn, Ryoji Miyazaki, Sixian Wu, Hiroyuki Tanaka, Yutaro S. Takahashi, Kaito Hosoda, Kunihito Yoshikaie, Yutetsu Kuruma, Hideki Shigematsu, Takaharu Mori, Tomoya Tsukazaki

## Abstract

Many antibiotics target essential cellular processes. To combat multidrug-resistant bacteria, new antibacterial strategies are needed. In the peptidoglycan biogenesis pathway in *Escherichia coli*, MurJ, the lipid II flippase, is an essential membrane protein. The 37-residue protein M from the *Levivirus* phage, known as Lys^M^ or Sgl^M^, targets MurJ and induces cell lysis; however, its molecular mechanism remains unclear. Here, we present the cryo-EM structure of the MurJ/Lys^M^ (JM) complex at 3.09 Å resolution, revealing that Lys^M^ interacts with the crevasse between TM2 and TM7 of MurJ, locking MurJ in an outward-facing conformation, with Lys^M^ acting like a wedge. Alanine-scanning mutagenesis and pull-down assays revealed key residues responsible for Lys^M^ function, and molecular dynamics simulations showed that Lys^M^ stabilizes MurJ’s outward-facing state. These findings demonstrate an unprecedented phage-derived mechanism for blocking lipid II transport, providing a structural framework for designing MurJ-targeted antimicrobial agents.

## Introduction

The emergence of multidrug-resistant bacteria seriously threatens public health, underscoring the urgent need for novel classes of antibiotics. In gram-negative bacteria, the peptidoglycan (PG) layer located between the inner and outer membranes is a fundamental structural component that maintains cell shape and enables adaptation to environmental changes^1,2^. Accordingly, many antibiotics, such as penicillin, target PG biogenesis to disrupt cell wall synthesis.

The PG layer primarily consists of glycan chains composed of repeating units of N-acetylglucosamine (GlcNAc) and N-acetylmuramic acid (MurNAc). Tetrapeptides attached to MurNAc are crosslinked with peptides from adjacent strands, forming a polymeric mesh structure in the periplasm (Fig. 1A). PG synthesis is initiated on the cytoplasmic side of the inner membrane. First, muramyl ligases (MurA–F) synthesize UDP-MurNAc-pentapeptide. This intermediate is then transferred to undecaprenyl phosphate by MraY, generating lipid I (Und-PP-MurNAc-pentapeptide). Subsequently, MurG catalyzes the formation of lipid II by adding UDP-GlcNAc to lipid I. Lipid II, initially facing the cytoplasm, is translocated across the membrane to the periplasmic side by the flippase MurJ^3–6^. Finally, lipid II is incorporated into the PG layer through a series of enzymatic steps, including polymerization.

**Figure 1.**
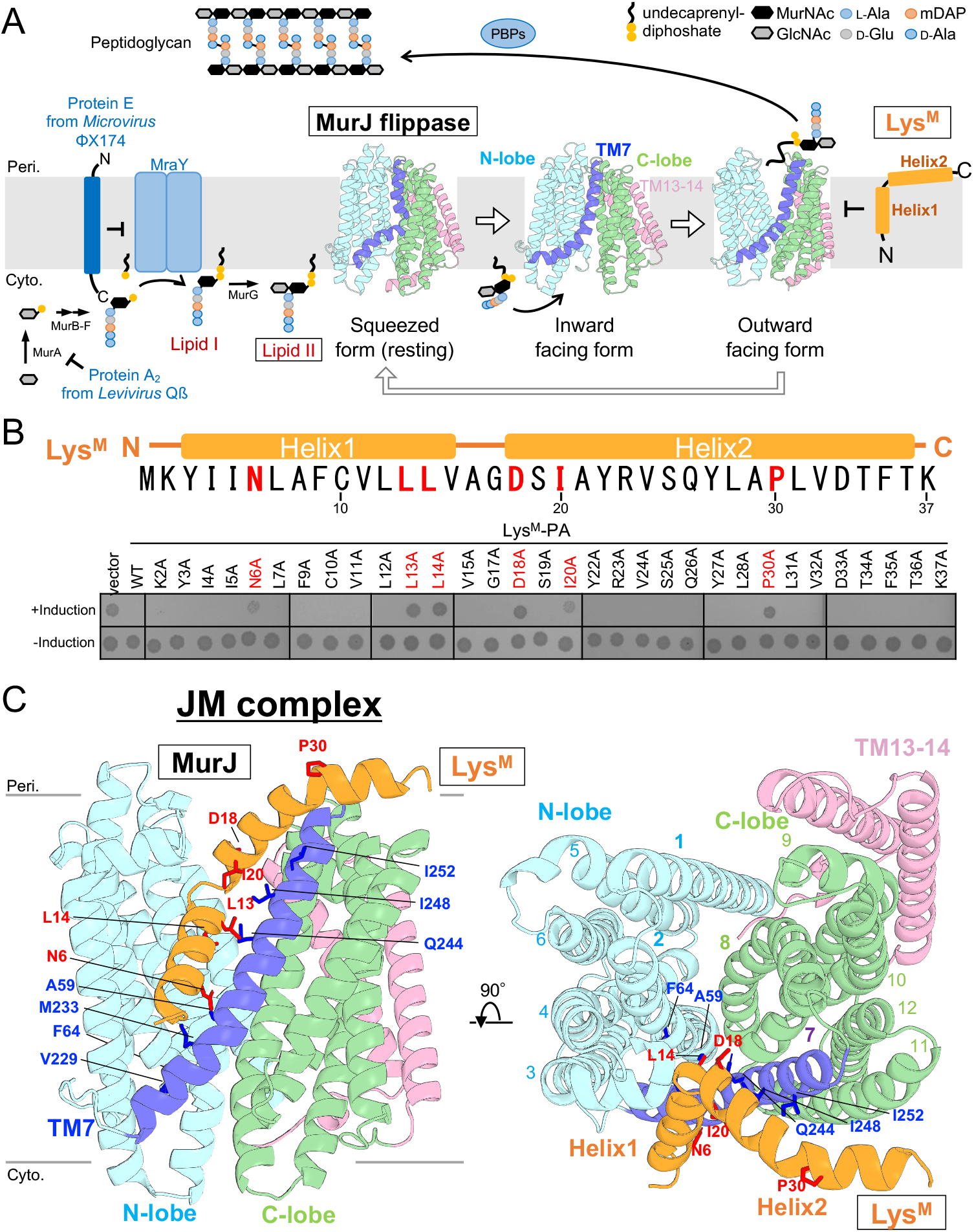
Structural basis for Lys^M^-mediated MurJ inhibition and cell lysis. (A) Overview of the peptidoglycan biosynthesis pathway and the conformational transitions of MurJ during lipid II flipping. Cartoon models of MurJ were generated based on available structures: the squeezed form of *E. coli* MurJ (PDB: 7WAG), the inward-closed form of *E. coli* MurJ (PDB: 6CC4), and the outward-facing form of *Thermosipho africanus* MurJ (PDB: 6NC9). The N-lobe (TM1–6), TM7 in the C-lobe, the remaining C-lobe region (TM8–12), and additional helices (TM13–14) are colored cyan, blue, green, and pink, respectively. The α-helical region of Lys^M^ is shown in orange. (B) Amino acid sequence and secondary structure of Lys^M^ with alanine-scanning mutagenesis. The six residues essential for cell lytic activity are highlighted in red. See also Figure 2A. (C) Cryo-EM structure of the JM (MurJ/Lys^M^) complex. MurJ and Lys^M^ are colored as in (A). Transmembrane helices are numbered in the right panel. Side chains of Lys^M^ residues essential for activity are shown as red sticks. Blue sticks represent the side chains of MurJ residues identified in suppressor mutants that confer resistance to Lys^M 14^.

Bacteriophages of the *Microviridae* and *Leviviridae* families are small single-stranded DNA and RNA phages that infect gram-negative bacteria and encode proteins that promote host cell lysis. Protein E from the microvirus ΦX174 and protein A2 from the *Levivirus* Qβ have been shown to inhibit MurA and MraY, respectively—both of which are essential enzymes in PG synthesis^7–11^ (Fig. 1A). A major recent breakthrough revealed the cryo-EM structure of the YES complex—composed of *Escherichia coli* MraY, protein E, and the host chaperone SlyD—which showed that protein E inhibits MraY activity by physically occupying its active site^12^. These phage-encoded proteins induce cell lysis by blocking PG biosynthetic enzymes without directly degrading the PG layer. Rumnieks and Tars identified *lys*^*M*^ (also known as *sgl*^*M*^) as a candidate lysis gene in the genome of *Levivirus* phage M, based on its expression in *E. coli*^*13*^. *lys*^*M*^ encodes a 37-residue protein, which is predicted to be a single membrane-spanning protein. Subsequently, Chamakura et al. cloned *lys*^*M*^ into a multicopy plasmid and analyzed the effects of Lys^M^ expression in *E. coli*, showing that Lys^M^ inhibits the lipid II flipping activity of *E. coli* MurJ, leading to cell lysis^14^.

MurJ is a membrane protein composed of 14 transmembrane helices (TMs), which are divided into three regions: the N-lobe (TM1–6), the C-lobe (TM7–12), and TM13–14. The N- and C-lobes form a V-shaped architecture that contains a central cavity for substrate binding. Structural and functional studies have suggested that MurJ transports lipid II from the cytoplasm to the periplasm via rocker-switch-like conformational changes between inward- and outward-facing states (Fig. 1B)^15–21^.

In the early stage of the transport cycle, lipid II enters the central cavity of the inward-facing form of MurJ. Upon binding, MurJ transitions to an inward-occluded form due to interactions between the lipid II headgroup and the internal cavity. The protein then undergoes a conformational change to an outward-facing form to release lipid II into the periplasm. After releasing lipid II, MurJ resets to the inward-facing state via a transient squeezed conformation to enable continuous transport. TM7, which extends from the C-lobe over the N-lobe, is proposed to play a central role in both inward-to-outward and outward-to-inward transitions (Fig. 1B)^16,17^. Accessibility experiments using MurJ mutants suggest that LysM locks MurJ in an outward-facing conformation by interacting with MurJ^14,22^. However, the precise mechanism by which Lys^M^ inhibits MurJ remains unclear.

Here, we performed comprehensive alanine-scanning mutagenesis of Lys^M^ and identified crucial residues required for its function. Furthermore, we determined the cryo-EM structure of the MurJ/Lys^M^ complex (termed the JM complex). Structural analysis revealed that Lys^M^ binds around TM7 of MurJ, a region involved in conformational transitions, and locks MurJ in an outward-facing conformation. These findings suggest that Lys^M^ functions as a conformational wedge, stabilizing MurJ and preventing its conformational cycling, thereby blocking lipid II transport—a process essential for peptidoglycan synthesis—and ultimately inducing cell lysis in *E. coli*. In addition, we discuss in detail the functional roles of several important residues of Lys^M^, based on structural and mutational analyses. These results provide a structural basis for the rational design of antimicrobial agents targeting MurJ-mediated lipid II transport.

## Results and Discussion

### Key amino acids required for Lys^M^ function

We first confirmed the cell lytic activity of Lys^M^, as previously reported ^13,14^. The *lysM* gene was cloned into pBAD33 under the control of an arabinose-inducible promoter and introduced into *E. coli* BL21(DE3) cells. Upon induction with L-arabinose, Lys^M^ expression led to a decrease in cell density, resulting in cell death and lysis (Fig. S1A). Lys^M^ derivatives fused to various C-terminal tags (His_10_-tag, PA-tag, or GFP-His_6_-tag) exhibited lytic activity comparable to that of untagged Lys^M^, indicating that these tags do not interfere with its function (Fig. S1A, B, Fig. S2B).

To identify residues critical for Lys^M^ function, we performed a comprehensive mutational analysis using alanine-substituted Lys^M^-PA mutants. Each non-alanine residue within the Lys^M^ region (residues 2–37) was individually substituted with alanine, resulting in a total of 32 Lys^M^-PA mutants. These mutants were then assessed for their cell lytic activity. Among them, six mutations—N6A, L13A, L14A, D18A, I20A, and P30A—failed to induce cell lysis and allowed normal cell growth even in the presence of arabinose (Fig. 1B, Fig. S2A), indicating a complete loss of function. To confirm that the loss of function was not due to reduced protein stability, we examined the accumulation of Lys^M^-GFP-His_6_ in *E. coli* cells. Expression of the *Bacillus subtilis* lipid II flippase Amj, which functions independently of MurJ^23^, allows lipid II transport in the presence of Lys^M^ in *E. coli*^*14,24*^. All Lys^M^-GFP-His_6_ variants, including the six loss-of-function mutants, accumulated to levels comparable to that of the wild-type upon Amj co-expression (Fig. S1E). These results suggest that N6, L13, L14, D18, I20, and P30 play essential roles in the lytic activity of Lys^M^.

### Structure of the MurJ/^LysM^ (JM) complex

We successfully co-purified Lys^M^-His_10_ and *E. coli* MurJ-3×FLAG from *E. coli* cells co-expressing Lys^M^-His_10_, MurJ-3×FLAG, and *B. subtilis* Amj, indicating that Lys^M^ directly interacts with MurJ. We reconstituted the MurJ/Lys^M^ complex into nanodiscs and performed cryo-EM single-particle analysis. The cryo-EM structure of the MurJ/Lys^M^ complex was determined at a resolution of 3.09 Å (Fig. 1C, Fig. S5, Table S1). The *E. coli* MurJ structure adopts an outward-facing form, distinct from previously reported inward-facing or “squeezed” forms^16,20^; an outward-facing conformation of *E. coli* MurJ has not been reported previously. The structure features a cavity opening toward the periplasm, formed by TM1, TM2, TM7, and TM8. Notably, an additional density not attributable to MurJ was observed in the cryo-EM map and assigned to Lys^M^. Structural modeling revealed that this density accommodates the full-length sequence of Lys^M^ (Fig. S5D).

The structure of the MurJ/Lys^M^ complex revealed that Lys^M^ consists of two helices: Helix 1 at the N-terminus and Helix 2 at the C-terminus (Fig. 1B, C). Although the TMHMM v2.0^25^ server predicted Lys^M^ to be a single membrane-spanning protein, the present structure shows that Lys^M^ is membrane-anchored. Helix 1 is deeply embedded in the lipid bilayer, whereas Helix 2 is oriented horizontally along the membrane interface (Fig. 1C). It is important to note that AlphaFold^26,27^ predictions did not reproduce MurJ/Lys^M^ complex structures resembling the cryo-EM structure. MurJ was consistently predicted in inward-facing forms, and the structural models of Lys^M^ were generated with low confidence (Fig. S3). Because structure prediction algorithms rely on existing PDB data, the experimentally determined outward-facing MurJ structure with Lys^M^ observed in this study might not be accurately predicted computationally. As discussed below, our structure captures a snapshot of Lys^M^ targeting MurJ and obstructing lipid II transport. We refer to this structure as the JM complex, named for the Lys<u>^M^</u>-mediated jamming of lipid II flipping by Mur<u>J</u>.

### C-terminal amphipathic helix of Lys^M^

To investigate the functional importance of Helix 2 at the C-terminus of Lys^M^, we constructed a series of C-terminally truncated mutants by introducing stop codons into the *lys*^*M*^ gene. Mutants retaining up to residues 1–35 and 1–36 exhibited cell lytic activity comparable to that of full-length LysM-His_10_. Mutants 1–33 and 1–34 showed slightly reduced activity but retained lytic function. In contrast, truncation mutants lacking residues beyond 1–32 completely lost lytic activity (Fig. S2C).

Structural analysis revealed that Helix 2 is an amphipathic helix aligned along the lipid bilayer surface (Fig. S2B). This amphipathic character likely contributes to the stable localization of Lys^M^ at the membrane interface. Notably, disruption of the hydrophobic–hydrophilic balance of Helix 2 may impair the proper membrane association of Lys^M^. Therefore, the amphipathic helix at the C-terminus could be important for both the function and membrane stability of Lys^M^.

### Lys^M^ locks MurJ in the outward-facing form

We examined how Lys^M^ interacts with MurJ in the JM complex structure. Helix 1 of Lys^M^ is positioned within a crevasse between TM2 and TM7 of MurJ. Notably, the loop connecting the two helices of Lys^M^ inserts into the lateral side of the MurJ cavity (Fig. 1C). To integrate structural and functional data, we mapped the six functionally important residues of Lys^M^ onto the JM complex structure. Five of these residues (N6, L13, L14, D18, and I20) are located at the Lys^M^–MurJ interface and directly contact MurJ (Fig. 1C).

In addition, we mapped the positions of seven MurJ mutations that confer resistance to Lys^M^, as identified in a genetic screen^14^. Almost all of the substituted residues are located at the Lys^M^–MurJ interface (Fig. 1C). These observations strongly support that the interaction between Lys^M^ and MurJ is critical for Lys^M^-mediated cell lysis. The JM complex structure is also consistent with a previous SCAM assay^14^, which demonstrated that MurJ adopts a conformation in which its central cavity becomes accessible from the periplasm in the presence of Lys^M^. Together, the cryo-EM structure and functional analyses suggest that Lys^M^ inhibits the conformational transition cycle of MurJ by strongly stabilizing it in the outward-facing state.

### Conformational transitions of *E. coli* MurJ

MurJ has been proposed to adopt multiple conformational states—inward-facing, inward-occluded, outward-facing, and squeezed—during lipid II transport^16,17^. With the outward-facing form of *E. coli* MurJ solved in this study, three distinct conformations (inward-facing, outward-facing, and squeezed) are now available, allowing a detailed discussion of its structural transitions (Fig. 2A). Among the transmembrane helices of MurJ, TM7 exhibits the most prominent conformational change. In the Lys^M^-bound outward-facing structure, TM7 adopts a straight conformation, whereas in the squeezed and inward-facing structures, TM7 bends around residue G239. The bending angle at G239 is approximately 60° in the squeezed state, ~40° in the inward-facing state, and only 14° in the JM structure (Fig. 2A). This comparison highlights TM7 as a key structural element involved in the conformational transitions of MurJ.

**Figure 2.**
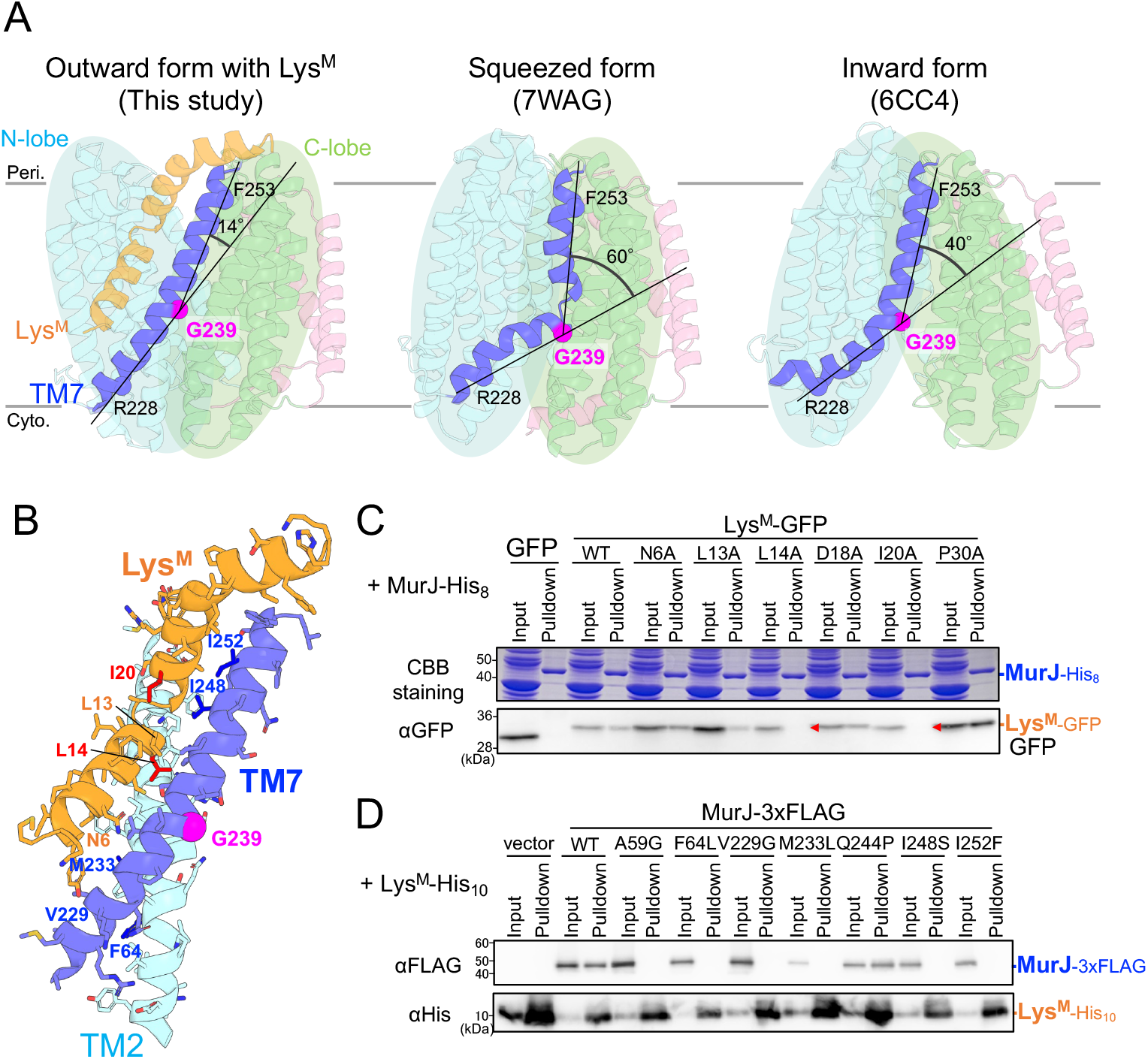
Lys^M^ targets a key region for conformational transitions in MurJ. (A) Structural comparison of *E. coli* MurJ in multiple conformations: inward-facing, squeezed, and outward-facing (bound to Lys^M^). The bending angle of TM7 at G239—defined by the Cα atoms of residues R228, G239, and F253—is indicated for each structure. (B) Interactions between TM2 and TM7 of MurJ and Lys^M^. TM2 and TM7 of MurJ and Lys^M^ are shown as cartoon models; the side chains are shown as sticks. Key residues involved in the Lys^M^–MurJ interaction in are mapped onto the model and colored red and blue. (C, D) Pull-down assays showing interactions between MurJ and Lys^M^ using various mutants, including Lys^M^-resistant mutants of MurJ and loss-of-function mutants of Lys^M^. See also Supplementary Figure 4.

### JM complex interaction changed by mutations

To investigate the binding ability of the six Lys^M^ mutants that lack lytic activity, we performed pull-down assays using *E. coli* cells co-expressing EcMurJ-His_8_, Lys^M^-GFP, and BsAmj. After Ni-NTA column chromatography, EcMurJ-His_8_ was detected through CBB staining, and the pull-down fractions were analyzed via immunoblotting with an anti-GFP antibody. Among the six mutants, L14A and I20A were undetectable in the pull-down fractions, indicating reduced binding to MurJ compared to wild-type Lys^M^ (Fig. 2C, Fig. S4A). In the cryo-EM structure, L14 and I20 are located at the interface between the N- and C-lobes of MurJ (Fig. 2B). These results demonstrate that interactions between MurJ and these residues are critical for lytic activity. Substitution of leucine and isoleucine with alanine shortens the side chains and likely weakens the hydrophobic interactions required for stable insertion into the MurJ interface.

We next examined the binding ability of MurJ mutants previously identified as Lys^M^-resistant. Pull-down assays were conducted using *E. coli* cells co-expressing EcMurJ-3×FLAG, Lys^M^-His_10_, and BsAmj. Both MurJ and Lys^M^ were detected via immunoblotting. Except for the Q244P mutant, all Lys^M^-resistant MurJ variants showed no detectable MurJ-FLAG signal in the pull-down fractions, indicating diminished interaction with Lys^M^ (Fig. 2D, Fig. S4B). These MurJ mutations, which impair Lys^M^-mediated lytic activity, appear to disrupt the physical interaction between the two proteins. All mutation sites were located around the Lys^M^–MurJ interface; in particular, M233, I248, and I252 in TM7 were found to make direct contact with LysM (Fig. 2B).

Because the lytic activity of Lys^M^ depends on its interaction with MurJ, these findings collectively indicate that direct physical binding to MurJ is essential for Lys^M^ function. Notably, the JM structure shows that Lys^M^ binds over the hinge residue G239 in TM7 of MurJ, locking TM7 in a straight conformation and thereby stabilizing the outward-facing state of MurJ.

The electrostatic surface of the JM complex revealed that outward-facing MurJ possesses a positively charged cavity open to the periplasm (Fig. 3). This cavity is critical for lipid II recognition, as it interacts with the phosphate groups of lipid II^15,28^. The side chains of Lys^M^ residues D18 and L14 extend into the cavity and make contact with the surface of MurJ (Fig. 3A). The negatively charged residue D18, located on the loop between Helix 1 and Helix 2 of Lys^M^, is likely positioned near the cavity through electrostatic interactions. In addition, alanine scanning and pull-down assays demonstrated that the D18A mutant retained binding ability to MurJ but completely lost lytic activity (Fig. S2A, Fig. 2C). These results suggest that although D18 is not essential for binding, it is important for stabilizing the outward-facing conformation of MurJ through electrostatic interaction. The side chain of L14, located at the tip of the loop, is surrounded by the side chains of L50, L54, V240, and Q244 of MurJ (Fig. 3B). This structural arrangement suggests that the L14 side chain plays a key role in forming the molecular wedge that locks MurJ in its outward-facing conformation.

**Figure 3.**
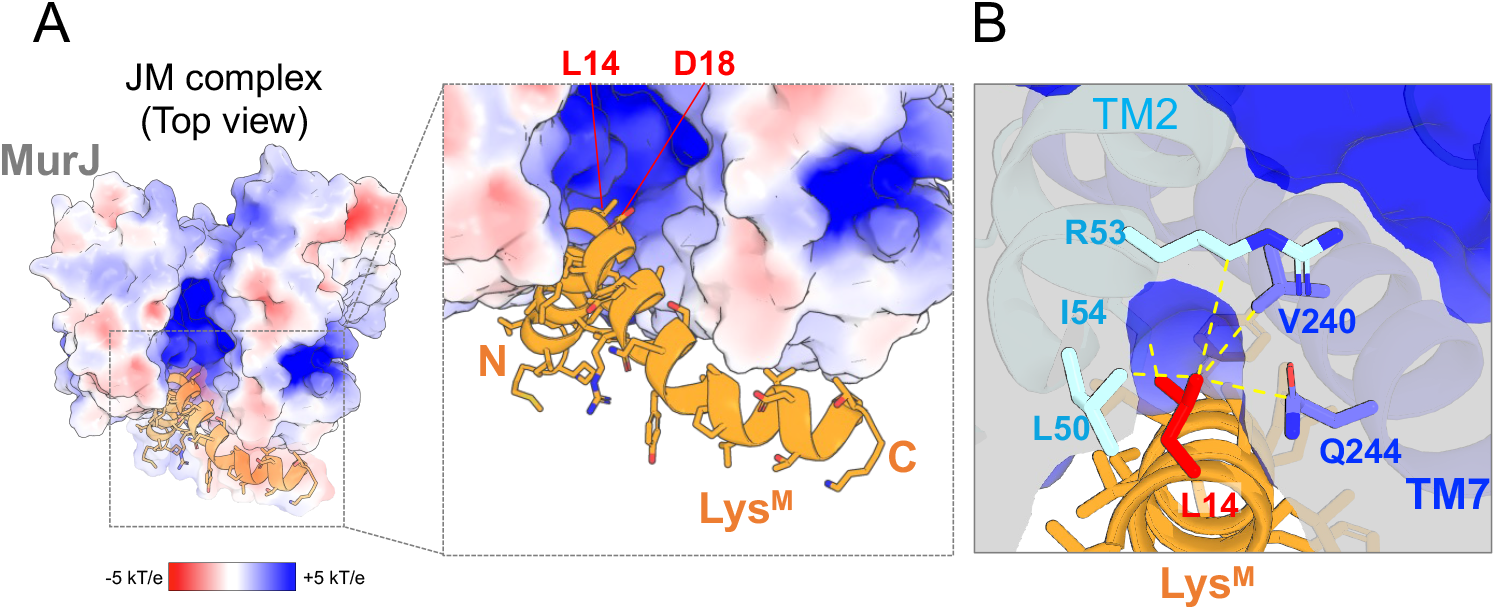
Electrostatic surface around the kink region of Lys^M^. (A) Top view of the JM complex. MurJ is shown as an electrostatic potential surface, colored from red (negative charge) to blue (positive charge). Lys^M^ is displayed in cartoon representation. (B) Close-up view of the region surrounding Lys^M^ residue L14, shown with a cutaway model. Dotted yellow lines indicate atomic distances (<5.0 Å) between the side chain of Lys^M^ L14 and the side chains of nearby MurJ residues.

### Molecular dynamics (MD) simulations of the JM complex

To gain insight into how Lys^M^ binding affects the structural dynamics of MurJ, we performed MD simulations of MurJ embedded in a membrane, both in the presence and absence of Lys^M^. In the MD simulations, the root-mean-square deviation (RMSD) of the JM complex remained low, indicating that the complex maintained structural stability throughout the simulation period (Fig. S6). In contrast, several MD simulations of outward-facing MurJ without Lys^M^ exhibited increases in RMSD values over time, suggesting that MurJ alone undergoes greater structural fluctuations compared to the JM complex. These *in silico* analyses support the idea that Lys^M^ stabilizes and locks MurJ in its outward-facing conformation.

To probe conformational flexibility in living cells, we introduced cysteine substitutions at specific residues and performed *in vivo* disulfide crosslinking experiments (Fig. S7). Disulfide bonds were observed between L14C, A16C, or S19C of Lys^M^, which retain their lytic activity, and I252C of MurJ, as well as between L14C of Lys^M^ and I245C of MurJ (Fig. S7B, C), indicating that TM7 of MurJ is positioned near the loop region of Lys^M^ in living cells. Interestingly, disulfide crosslinked products were also detected between residues that do not directly contact each other in the cryo-EM structure, such as L14 of Lys^M^ and I252 of MurJ (Fig. S7A, B). These observations suggest that Helix 1 of Lys^M^ initially resides in a shallower position within the membrane before forming a tightly bound complex with MurJ.

Moreover, MD simulations of Lys^M^ alone revealed that the protein adopts either a single membrane-spanning configuration, as predicted by TMHMM, or a bent conformation within the membrane, in which its amphipathic Helix 2 lies horizontally along the membrane interface (Fig. S6). These simulations indicate that Lys^M^ undergoes dynamic conformational changes within the membrane environment. This conformational flexibility may be restricted upon binding to its partner protein MurJ, thereby contributing to the enhanced structural stability of the JM complex.

### Model of Lys^M^-induced block on lipid II transport via MurJ

Based on the structural analyses presented in this study, we updated the working model of MurJ-mediated lipid II transport and proposed a mechanism by which Lys^M^ inhibits MurJ function (Fig. 4). The cryo-EM structure of the JM complex revealed an outward-facing conformation of *E. coli* MurJ, which complements previously reported inward-facing and squeezed conformations, completing the structural cycle of *E. coli* MurJ during lipid II transport. In this model, MurJ initially adopts an inward-facing conformation open to the cytoplasm, where it binds lipid II. It then transitions to an outward-facing conformation to release lipid II into the periplasm and then returns to a squeezed state, which likely represents a resting conformation. A key structural element in this cycle is TM7, which undergoes bending and straightening through a hinge centered at residue G239.

**Figure 4.**
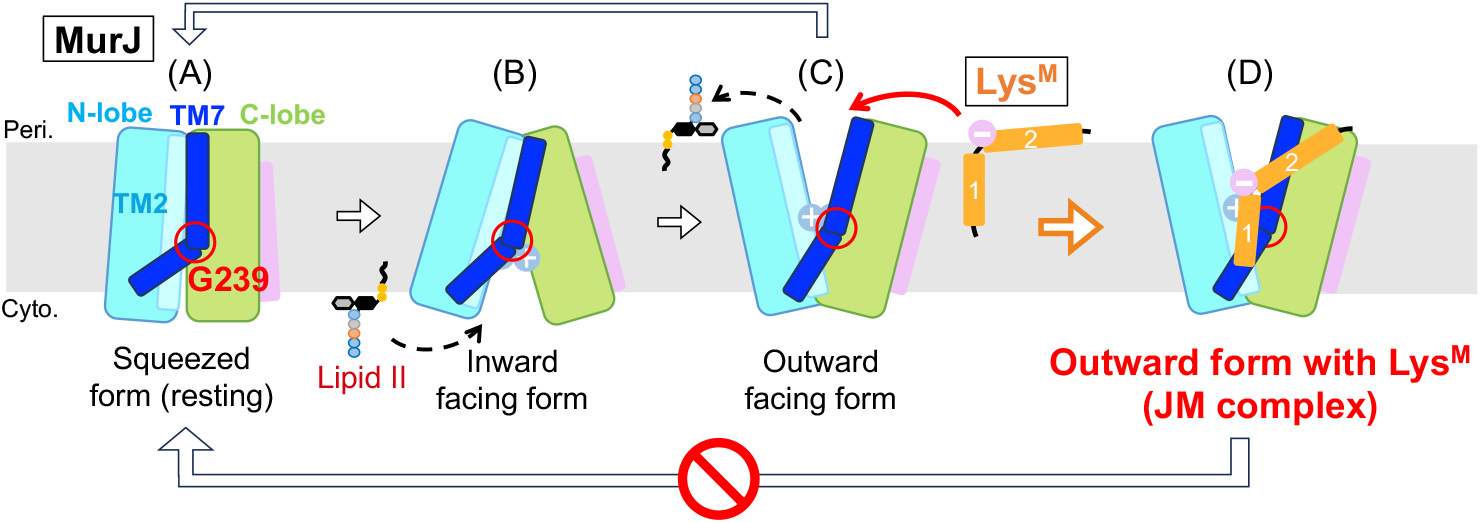
Working model for Lys^M^-mediated inhibition of MurJ. (A–C) Schematic illustration of the lipid II transport cycle mediated by MurJ. (A) The squeezed form represents the resting state of MurJ. (B) The inward-facing form captures lipid II on the cytoplasmic side. (C) The outward-facing form opens toward the periplasm, allowing the release of lipid II. After release, MurJ returns to the squeezed state (A) to complete the transport cycle. (D) Lys^M^ binds to MurJ in the outward-facing conformation. Lys^M^ restricts the flexibility of residue G239 in TM7, thereby limiting its bending. Lys^M^ functions as a conformational wedge, inhibiting the rocker-switch-like transition of MurJ.

Lys^M^ inhibits MurJ by locking it in the outward-facing conformation. Insertion of the Lys^M^ loop into the groove between TM2 and TM7 of MurJ prevents further conformational transitions and halts lipid II transport. The stabilization of the JM complex is supported by three structural features: (i) the amphipathic Helix 2 of Lys^M^ positioned along the membrane surface, (ii) hydrophobic interactions between Helix 1 of Lys^M^ and MurJ, and (iii) electrostatic interactions between the negatively charged Lys^M^ loop and the positively charged cavity of MurJ. Thus, Lys^M^ functions as a molecular wedge against MurJ, mechanically jamming the lipid II flippase cycle and blocking a critical step in peptidoglycan biosynthesis.

Understanding the mechanism of ssRNA phage lysis proteins, such as Lys^M^, may aid in the development of novel antibiotics, including variants tailored to specific bacterial hosts. Furthermore, the small size and high specificity of Lys^M^ make it a promising candidate for applications in synthetic biology, such as regulating cell growth.

## Material and Methods

### Bacterial Strains and Plasmids

The *E. coli* strains and plasmids used in this study are listed in Table S2. Details of the plasmid used are described in the “Plasmid Construction” section.

### Plasmid Construction

A DNA fragment containing the *lys*^*M*^ gene and its Shine-Dalgarno sequence (5’-CTCGGTACCCGGGGATCCTCTAGGAGGTTTAAATTATGAAATATATAATAAATTTAGCATTCTGCGTTCTCTTACTGGTTGCGGGGGACTCGATAGCATATCGAGTCTCGCAATACCTGGCGCCTTTGGTGGATACCTTCACCAAGTAAAGAGTCGACCTGCAGGCATGC-3’) and XbaI-digested pBAD33 were changed into a plasmid by Gibson assembly^29^. The resulting plasmid was named pKK568. A DNA sequence (5’-GGAGTCGCCATGCCCGGAGCCGAGGATGATGTCGTC-3’) encoding the PA tag was inserted before the stop codon of *lys*^*M*^ by site-directed mutagenesis, and the resulting plasmid was named pKG74. The plasmid pKG191 (pBAD18 encoding *lys*^*M*^*-his*_*10*_) was constructed in a similar way to pKG74. For pKG126, the *GFP-HIS*_*6*_ gene was PCR-amplified from pCGFP-BC^30^. The PCR product was cloned into SalI-PstI-digested pKK568 using In-Fusion HD Cloning Kit (Takara Bio). The plasmids with mutations were constructed using site-directed mutagenesis.

For pKG537 (pSTV28 encoding *ecmurJ-3xflag* [DYKDHDGDYKDHDIDYKDDDDK]), the *E. coli murJ* fragment was PCR-amplified from pKG65 using specific primers. The PCR product was cloned into the corresponding sites in pSTV28 using the In-Fusion HD Cloning Kit (Takara Bio). The plasmids with mutations were constructed using site-directed mutagenesis.

For pRM1071, a *Bacillus subtilis Amj* fragment was PCR-amplified from the genome of *B. subtilis* subsp. *subtilis* JCM 1465T (RIKEN BRC) using specific primers (Bs_amj-F, 5’-GCGCGAATTCATTGGAGGAAGAATAACGTGC-3’; Bs_amj-R, 5’-GCGCAAGCTTTTAAAACCACTTTGTCAGCC-3’). The PCR product was digested with EcoRI and HindIII and ligated to the corresponding sites in pTWV228 (Takara Bio). The resulting plasmid was named pRM1071.

For pRM1210, a DNA fragment containing the *B. subtilis amj* was PCR-amplified from pRM1071 using specific primers (Bs_amj-F2, 5’-CAGGAAACAGCCATGCATGTCATTACAACACAAG-3’; Bs_amj-R2, 5’-GGCCAGTGCCAAGCTTTTAAAAC-3’) and cloned into NcoI-HindIII-digested pHM1552 (Miyazaki et al., 2022) using an In-Fusion HD Cloning Kit (Takara bio).

### Media and Bacterial Cultures

*E. coli* cells were grown at 37°C in LB medium (Nacalai Tesque). 100 μg/mL of ampicillin (Amp), 20 μg/mL of chloramphenicol (Cm), and 50 μg/mL of spectinomycin (Spc) were added as appropriate for the growth of plasmid-bearing cells and selection of transformant. Bacterial growth was monitored with a Mini photo 518R (660 nm; TAITEC Co., Japan).

### Lysis assay

*E. coli* BL21(DE3) cells carrying a plasmid encoding Lys^M^ derivatives were grown overnight in LB medium supplemented with the appropriate antibiotics. Cultures were diluted 1:10,000 in PBS and spotted onto LB agar plates supplemented with 0.2% (w/v) L-arabinose and the necessary antibiotics. The plates were incubated at 37°C for 22 h and subsequently imaged.

### Purification of the MurJ/Lys^M^ complex

*E. coli* BL21 Δ*recA*(DE3) harboring plasmids pKG537, pKG191, and pRM1210, encoding MA-*Ec*MurJ_2-511_ (from *E. coli* JCM20135)-3xFLAG, Lys^M^-His_10_, and *Bs*Amj, respectively, was cultured in LB medium (Lennox, Nacalai) supplemented with 100 μg/mL Amp, 20 μg/mL Cm, and 50 μg/mL Spc at 37°C. Upon reaching an OD_660_ of 0.7–0.8, protein expression was induced by adding 1 mM isopropyl β-D-thiogalactopyranoside (IPTG) and 0.2% (w/v) L-arabinose. Cultures were then incubated at 22°C for 18 h. Harvested cells were suspended in buffer (20 mM Tris-HCl pH 8.0, 150 mM NaCl, and 0.1 mM phenylmethylsulfonyl fluoride [PMSF]) and lysed using an M-110EH Microfluidizer (Microfluidics) at 15,000 psi (three passes). The cell lysate was centrifuged at 12,000 rpm for 30 min (Hitachi R13A rotor), and the supernatant was ultracentrifuged at 40,000 rpm for 60 min (Beckman 45Ti rotor). The pellet was resuspended in solubilization buffer (20 mM Tris-HCl, pH 8.0, 300 mM NaCl, 2% [w/v] n-dodecyl β-maltoside [DDM], 0.02% lauryl maltose neopentyl glycol [LMNG], 0.002% cholesterol hydrogen succinate [CHS] [w/v], 20 mM imidazole-HCl, pH 8.0, and 0.1 mM PMSF) and rotated for 60 min at 4°C. Following ultracentrifugation for 30 min, the supernatant was incubated with Ni-NTA agarose resin (Qiagen) for 60 min at 4°C. The resin was washed, and bound proteins were eluted with an elution buffer (20 mM Tris-HCl, pH 8.0, 300 mM NaCl, 0.02% LMNG, 0.002% CHS, 300 mM imidazole-HCl, pH 8.0, and 0.1 mM PMSF). Eluted fractions containing MurJ-3xFLAG and Lys^M^-His_10_ were collected and concentrated using an Amicon Ultra 50 K NMWL (Merck Millipore). After ultracentrifugation (40,000 rpm for 30 min, himac S55A2 rotor), the concentrated sample was then applied to a Superose 6 increase 10/300 GL column (Cytiva) equilibrated with buffer (20 mM Tris-HCl, pH 8.0, 300 mM NaCl, 0.005% LMNG, 0.0005% CHS, and 0.1 mM PMSF). Fractions containing the target proteins were collected and concentrated to ~20 mg/mL using an Amicon Ultra 50 K NMWL.

### Nanodisc-reconstitution of the MurJ/Lys^M^ complex

The purified MurJ/Lys^M^ complex was mixed with MSP1E3D1^31^ and *E. coli* phospholipids (Avanti) at a molar ratio of 1.4:1:120 in buffer (20 mM Tris-HCl, pH 8.0, 300 mM NaCl, 0.005% LMNG, 0.0005% CHS, and 0.1 mM PMSF). The mixture was incubated at 4°C for 1 h. After addition of Bio-Beads SM2 (Bio-Rad), the mixture was gently rotated overnight at 4°C to initiate detergent removal and nanodisc formation. Next, the mixture was filtered using Centrifugal Filters PVDF 0.22 μm (Millipore). The filtrate was then ultracentrifuged at 45,000 rpm for 30 min (himac S55A2 rotor). The resulting supernatant was applied to a Superdex 200 10/300 GL column pre-equilibrated with buffer (20 mM Tris-HCl, pH 8.0, 300 mM NaCl, and 0.1 mM PMSF). Fractions were analyzed by SDS-PAGE and native PAGE to confirm nanodisc reconstitution. Fractions containing the MurJ/Lys^M^-reconstituted nanodiscs were pooled and concentrated using an Amicon Ultra 50 K NMWL (Merck Millipore).

### Cryo-EM data collection and processing

Quantifoil holey carbon grids (Cu R1.2/R1.3, 300 mesh) were glow-discharged at 7 Pa and 10 mA for 10 s using a JEC-3000FC sputter coater (JEOL) before sample application. A 3 μL aliquot of 2.8 mg/mL of the MurJ/Lys^M^ complex-reconstituted nanodiscs was applied onto the grids, blotted for 3 s at 100% humidity, 8°C, and blot force 10, and plunged into liquid ethane using a Vitrobot Mark IV (Thermo Fisher Scientific).

The cryo-EM dataset was collected on a CRYO ARM 300 transmission electron microscope (JEOL) operated at an accelerating 300 kV, equipped with a cold-field emission gun, an in-column Omega-type energy filter, and a Gatan K3 direct electron detector (Gatan) at SPring-8. Images were collected at a nominal magnification of 60,000×, corresponding to a calibrated pixel size of 0.752 Å/pixel, 50 frames per image with a total exposure dose of 50 e^−^/Å^2^ using a SerialEM^32^.

The data processing was performed using CryoSPARC v4.5.3 software^33^. Also see the cryo-EM data-processing workflow for the MurJ/Lys^M^ complex in Figure S5A. A total of 7,973 movies were aligned using patchmotion correction, and the contrast transfer function (CTF) parameters were estimated using the Patch CTF estimation. The particles were automatically picked using the Blob picker in CryoSPARC among 200 of the 7,973 micrographs. Particles were extracted with a downsampling via Fourier cropping to a pixel size of 3.008 Å and subjected to 2D classification, generating 2D class averages. Using these 2D class averages as templates, particles were re-picked the Template Picker from 7,973 micrographs, yielding 2,994,327 particles. The repicked particles were extracted with a downsampling via Fourier cropping to the pixel size of 1.504 Å and subjected to two rounds of 2D classification (K = 200, 200). After 2D classification, approximately 1,000,000 high-quality particles were selected and re-extracted to 0.752 Å/pixel. These selected particles were subjected to ab initio 3D reconstruction (K = 4), generating initial 3D models. The 2,006,751 particles were then subjected to three rounds of heterogeneous refinement (K = 4, 4, 4) using the initial models as references, removing junk particles. The best 3D class, containing 317,838 particles, was re-extracted to 0.752 Å/pixel and refined using Non-Uniform (NU) refinement. These particles were subjected to reference-based motion correction and then subjected to Non-Uniform (NU) refinement, yielding a 3D cryo-EM map with an estimated overall resolution of 3.09 Å. Local resolution was estimated using the Local Resolution Estimation job in CryoSPARC.

### Model building and refinement

An *Ec*MurJ outward homology model was generated using the *Thermosipho africanus* (*Ta*MurJ) outward structure (PDB ID: 6NC9) as a template via the SWISS-MODEL Workspace (https://swissmodel.expasy.org/). The predicted Lys^M^ structure was obtained using AlphaFold3^27^. The resulting models were docked into the cryo-EM map using UCSF Chimera X. Manual model adjustments were performed in Coot^34^, and further refinement was conducted using the phenix.real_space_refine in PHENIX^35^. Data processing and refinement statistics are provided in Figure S5 and Table S1. The final molecular model and cryo-EM map were visualized using PyMOL (https://pymol.org/) or UCSF ChimeraX^36^.

### Pull-down assay

*E. coli* BL21 Δ*recA*(DE3) cells harboring plasmids encoding EcMurJ, Lys^M^ derivatives, and BsAmj were cultured in LB medium (Lennox, Nacalai) supplemented with 100 μg/ml Amp, 20 μg/mL Cm, and 50 μg/mL Spc at 37°C. When the culture reached an OD_600_ of 0.7–0.8, protein expression was induced by 1 mM isopropyl β-D-thiogalactopyranoside and 0.2% (w/v) L-arabinose. The cultures were then incubated at 25°C for 18 h. Cells were harvested and suspended in lysis buffer (20 mM Tris-HCl pH 8.0, 150 mM NaCl) and disrupted by sonication. The lysate was centrifuged at 12,000 rpm for 30 min using a Hitachi R13A rotor to remove debris and the supernatant was ultracentrifuged at 40,000 rpm for 60 min (Beckman 45Ti rotor) to isolate membrane fractions. The membrane pellets were resuspended in solubilization buffer (20 mM Tris-HCl pH 8.0, 300 mM NaCl, 2% DDM, 20 mM imidazole-HCl, pH 8.0) and incubated with gentle rotation for 60 min at 4°C. Following solubilization, the mixture was ultracentrifuged for 30 min, and the supernatant was incubated with 100 μl of Ni-NTA agarose resin (Qiagen) pre-equilibrated with wash buffer (20 mM Tris-HCl, pH 8.0, 300 mM NaCl, 0.1% DDM, 20 mM imidazole-HCl, pH 8.0) for 1 h at 4°C. After washing with the same buffer, bound proteins were eluted with elution buffer (20 mM Tris-HCl, pH 8.0, 300 mM NaCl, 0.1% DDM, 300 mM imidazole-HCl, pH 8.0). The eluted fractions were analyzed by immunoblotting.

### Immunoblotting analysis

Protein samples in SDS sample buffer (62.5 mM Tris-HCl pH 8.0, 2% SDS, 10% glycerol, 5% β-mercaptoethanol) were separated by SDS-PAGE using a 10% Laemmli gel. For analysis of Lys^M^ derivatives, proteins were resolved using a 15% Bis-Tris gel with MES running buffer, as described previously^37^. Following electrophoresis, proteins were transferred onto PVDF membranes (Merck Millipore). The membranes were blocked with skim milk and probed with one of the primary antibodies: anti-His (1:5,000 dilution, MBL #PM032), anti-GFP (1:5,000 dilution, MBL #598), or anti-FLAG M2 (1:50,000 dilution, Sigma-Aldrich #F1804). After washing with PBST, membranes were incubated with HRP-conjugated secondary antibodies: goat anti-rabbit IgG (1:3,000 dilution, BIO-RAD #170-6515) or anti-Mouse IgG (1:5,000 dilution, MBL #330). Protein bands were visualized using Chemi-Lumi One (Nacalai Tesque) and imaged with FUSION Solo S (VILBER).

### Lys^M^ x MurJ disulfide crosslinking

*E. coli* BL21Δ *recA* (DE3) carrying a combination of plasmids encoding a Cys-introduced mutant of MurJ-3xFLAG and Lys^M^-His_10_ were cultured in LB medium supplemented with 100 μg/ml Amp, 20 μg/mL Cm, and 50 μg/mL Spc. Protein expression was induced by adding 0.2% arabinose and 0.2 mM IPTG, and cultures were incubated at 30°C for 4 h. The harvested cells were washed, and resuspended in 20 mM Tris-HCl (pH 8.0). The cell suspensions were supplemented with 1 mM Cu(1,10-phenanthroline)_3_ solution and incubated at 37°C for 5 min. After that, the solution was supplemented with 15 mM EDTA. Total cellular proteins were precipitated with 5% TCA, washed with acetone, and suspended in SDS sample buffer containing 12.5 mM NEM to block free-SH groups of Cys residues. Half of the samples were mixed with 10% 2-mercaptoethanol for reducing and non-reducing analyses. Proteins were separated by 10% Laemmli SDS-PAGE and analyzed by immunoblotting using anti-His and anti-FLAG antibodies, as described above.

### MD simulations of MurJ/LysM, MurJ, and LysM in membranes

All-atom molecular dynamics (MD) simulations were performed for three systems: the MurJ/LysM complex, MurJ alone, and LysM alone. Each system was embedded in a mixed POPE/POPG lipid bilayer and solvated in a 150 mM NaCl solution. The initial structure of the protein-membrane complex was constructed using the CHARMM-GUI *membrane builder*^*38*^. The system sizes were as follows: 128 × 128 × 106 Å^3^ for the MurJ/LysM complex and MurJ alone (~167,000 atoms), and 100 × 100 × 94 Å^3^ for LysM alone (~91,000 atoms). The CHARMM36m force field was used for both proteins and lipids^39^. The system was equilibrated using 10,000-steps energy minimization, followed by equilibration in the *NVT* and *NPT* ensembles for 4.4 ns using the positional restraints on the proteins and lipids. Subsequently, production runs were carried out in the *NPT* ensemble at 303.15 K and 1 atm using a 2.5 fs time step, for 225 ns for both the MurJ/LysM complex and MurJ alone, and for 690 ns (Run1) and 667.5 ns (Run2) for LysM alone. The RESPA integrator was used with the Bussi thermostat and barostat for temperature and pressure control^40^. Long-range electrostatic interactions were treated with the particle-mesh Ewald method^41^. All simulations were performed using GENESIS 2.1.5^42^.

## Acknowledgments

We thank Kayo Abe for secretarial assistance and the scientists of SPring-8 Structural Biology Beamlines for helping with data collection. The cryo-EM experiments were performed at SPring-8 with the approval of the Japan Synchrotron Radiation Research Institute (Proposal Nos. 2024A2742, 2024A2759). This work was supported by the JSPS/MEXT KAKENHI (Grant No. JP25K18422, JP23K14146 to H.K., JP21H05156 to Y.K., Grant Nos. JP25K00267, JP24KK0138, JP22K15061, JP22H05567 to R.M., Grant Nos. JP21H05157, JP24K03035, 25H01329 to T.M., and Grant Nos. JP25K02226, JP22H02567, JP22H02586, JP21H05155, JP21H05153, JP21KK0125 to T.T.); private research foundations (The Chemo-Sero-Therapeutic Research Institute, ONO Medical Research Foundation, and Takeda Science Foundation to T.T., and the Institute for Fermentation (Osaka) to H.K., R.M. and T.T.); and JST SPRING (Grant No. JPMJSP2140 to Y.S.T). This research was partially supported by Platform Project for Supporting Drug Discovery and Life Science Research (Basis for Supporting Innovative Drug Discovery and Life Science Research (BINDS)) from AMED under Grant Number JP24ama121001. MEXT as “Program for Promoting Researches on the Supercomputer Fugaku” (Development and application of large-scale simulation-based inferences for biomolecules JPMXP1020230119 to T.M.), HPCI project (hp230209, hp240215, hp240277 to T.M.), and Research Center for Computational Science, Okazaki, Japan (Project: 25-IMS-C151 to T.M.). We thank GeneFrontier Corp. for providing the PURE system for trial use in the purification of Lys^M^. We gratefully acknowledge Nara Institute of Science and Technology for access to its facilities on weekends and public holidays, enabling part of this work as voluntary, unpaid research.

## Author contributions

H.K. and T.T. conceptualized the study. H.K., N.L., R.M., S.W., H.T., Y.K., and T.T. performed the biological analyses. H.K., Y.S.T., K.H., K.Y., H.S., and T.T. determined the structure. T.M. performed the MD simulations. H.K. and T.T. wrote the manuscript. H.K., T.M., and T.T. supervised the study.

## Data availability

The coordinates and the electron potential maps for the JM complex structure have been deposited in PDB and EMDB under accession codes PDB: 9UKV and EMBL-EBI: 64256, respectively.

