## Supplemental Figures and Tables for "Phage lysis protein Lys^M^ acts as a wedge to block MurJ conformational changes"

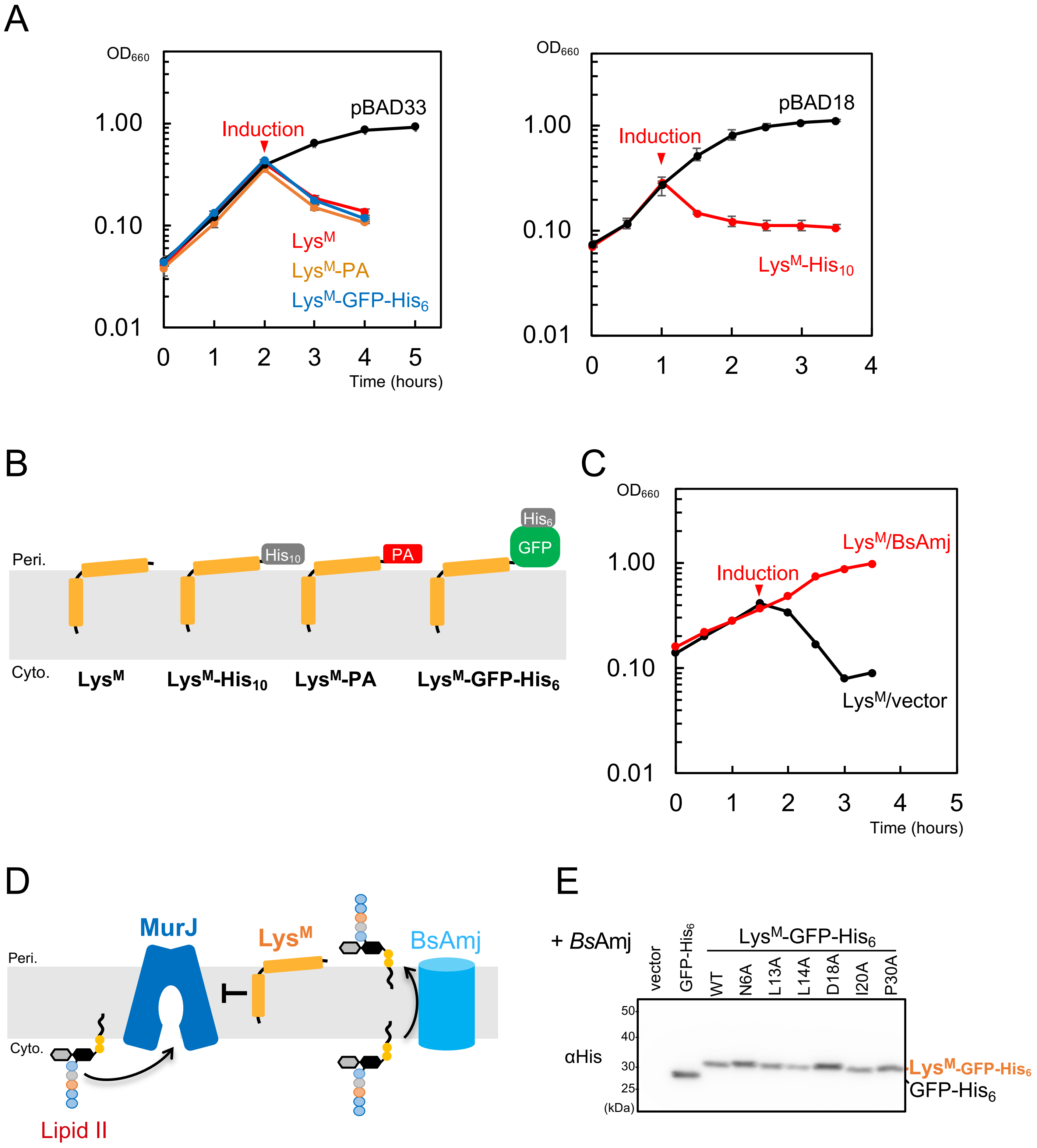


**Supplementary Figure 1. *E. coli* cell lysis by Lys$^{M}$**

(A) *E. coli* lysis induced by Lys$^{M}$ fused to His-tag, PA-tag, or GFP-His$_6$. Growth curves of *E. coli* cells harboring pBAD33/18 (vector) or plasmids encoding tagged versions of *lysM* are shown. Cells were grown in LB medium for 2 h and treated with 0.2% (w/v) L-arabinose at the 2-h time point. $OD_{660}$ was monitored. (B) Schematic representation of the Lys$^{M}$ derivatives used in this study. (C) Suppression of Lys$^{M}$-induced *E. coli* cell lysis by BsAmj. Growth curves of cells expressing untagged Lys$^{M}$ along with either pTWV228 (vector) or a plasmid encoding BsAmj. Cells were grown in LB medium for 1.5 h and induced with 0.2% (w/v) L-arabinose and 0.2 mM IPTG at 1.5 h. (D) Schematic illustrating BsAmj-mediated complementation of Lys$^{M}$-induced inhibition of MurJ-mediated lipid II transport. (E) Accumulation of Lys$^{M}$-GFP-His$_6$ in transformants shown in Fig. 2C. After a 2-h induction with L-arabinose and IPTG, total cell lysates were acid-precipitated and analyzed using SDS-PAGE followed by immunoblotting with an anti-His antibody.

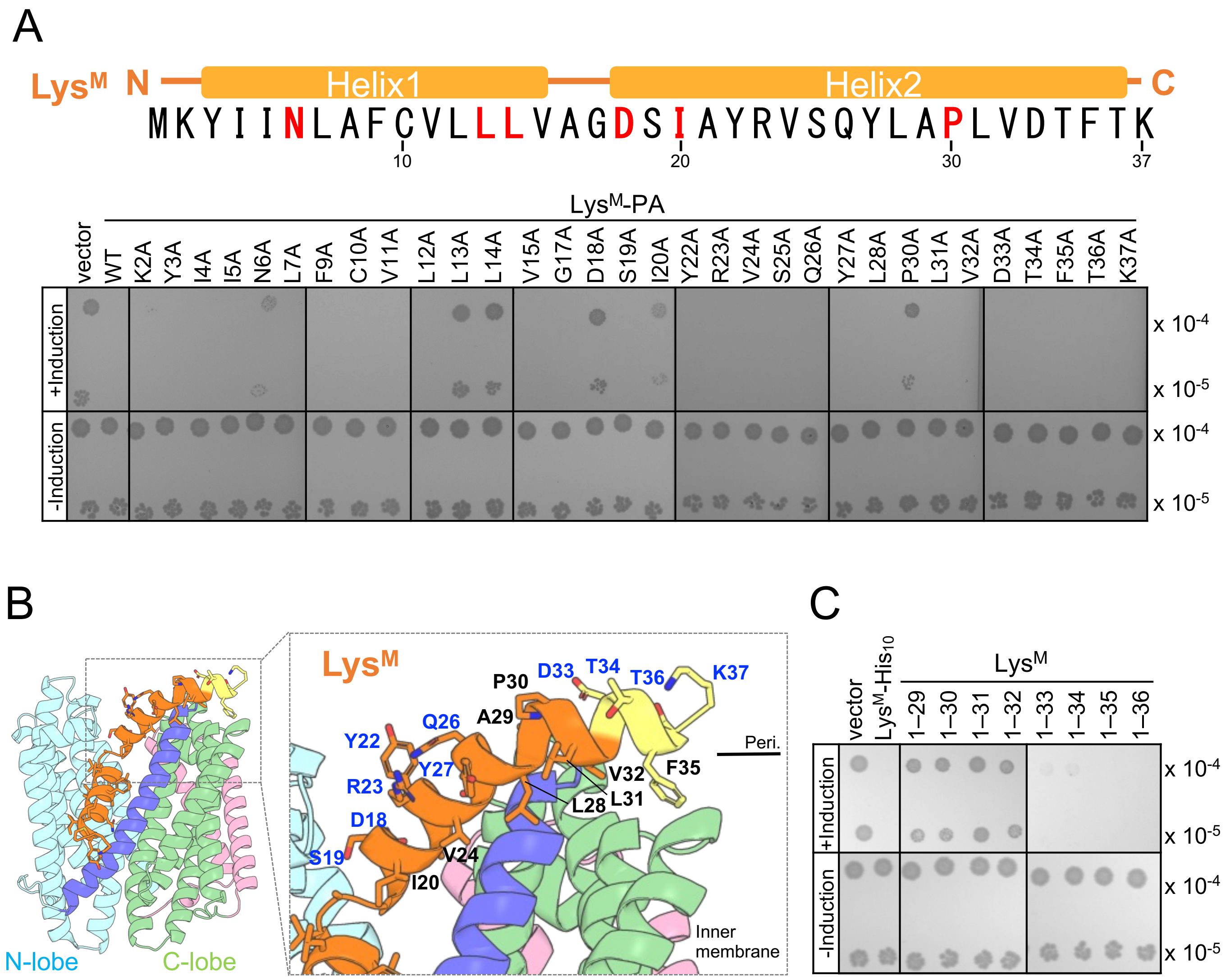


**Supplementary Figure 2. Essential residues for Lys$^{M}$ function**

(A) Effects of alanine-substituted Lys$^{M}$ mutants on *E. coli* growth. Amino acid sequence and secondary structure of Lys$^{M}$ (top). The six amino acid residues highlighted in red are identified as essential for cell lytic activity through alanine-scanning mutagenesis. *E. coli* cells harboring pBAD33 (vector) or plasmids encoding Lys$^{M}$-PA (or its mutants) were grown on LB agar plates with or without 0.2% (w/v) L-arabinose. (B) Helix 2 of Lys$^{M}$. Helix 2 exhibits an amphipathic α-helix. Side chains of Lys$^{M}$ residues are shown as sticks. (C) Cell lytic activity of C-terminal truncated Lys$^{M}$ mutants. *E. coli* cells harboring pBAD18 (vector) or plasmids encoding Lys$^{M}$ (or its mutants) were grown on LB agar plates with or without 0.2% (w/v) L-arabinose.

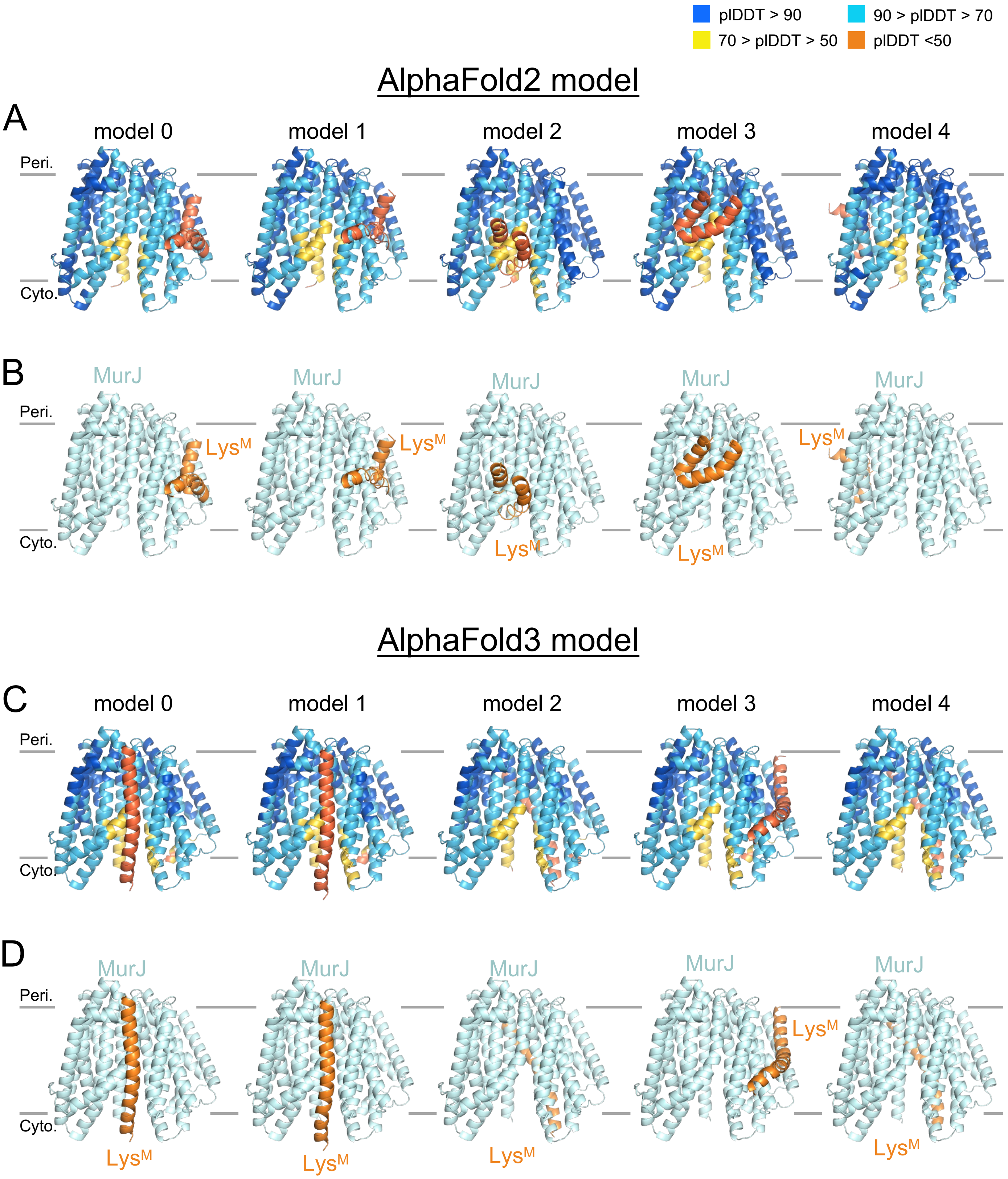


**Supplementary Figure 3. AlphaFold-predicted models of MurJ/Lys$^{M}$ complex.**
MurJ/Lys$^{M}$ models were generated using AlphaFold2 and AlphaFold3. pLDDT scores indicate structural accuracy. (A, C) Predicted models are color-coded based on per-residue pLDDT scores. (B, D) MurJ and Lys$^{M}$ are colored pale blue and orange, respectively.

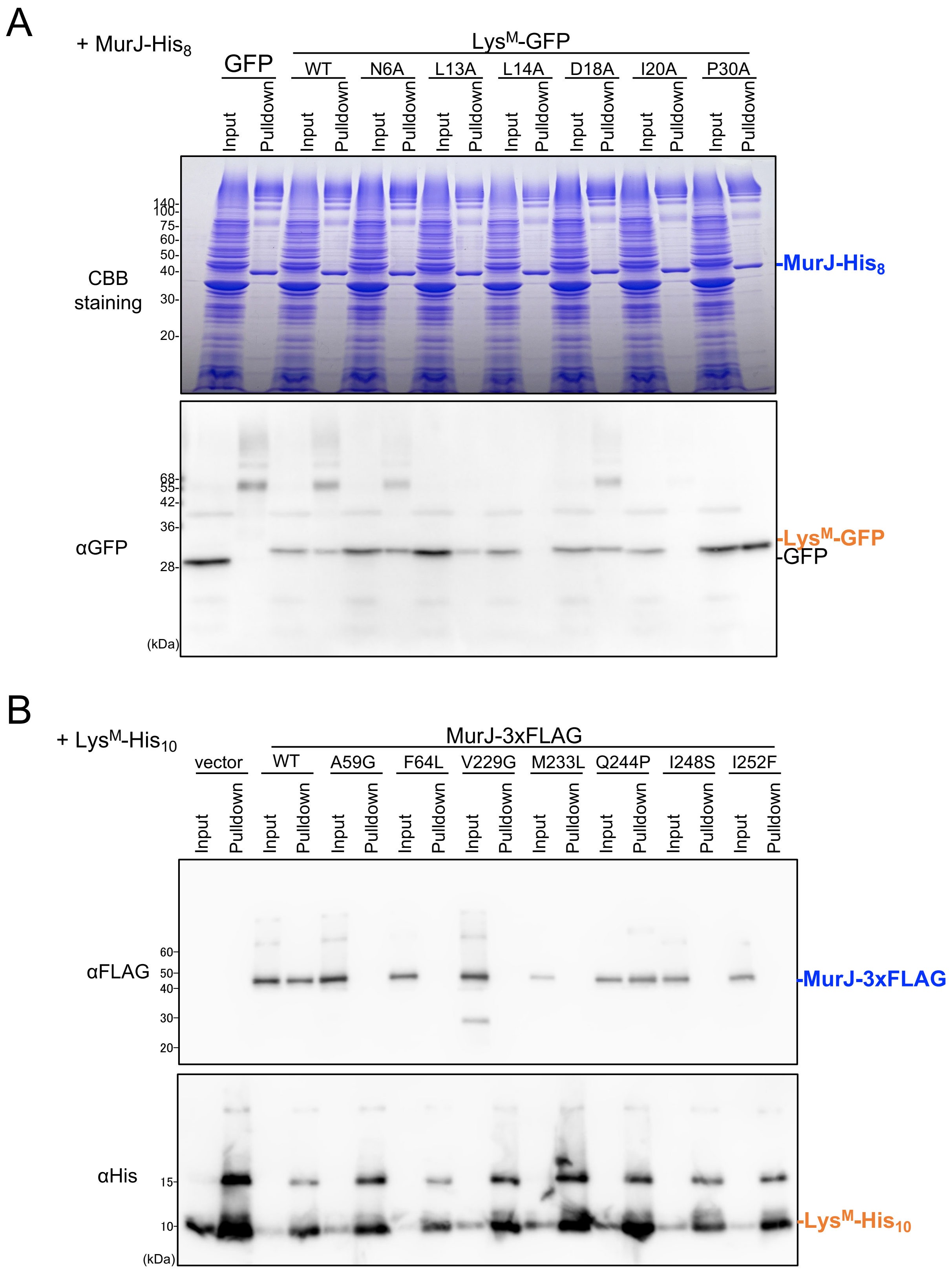


**Supplementary Figure 4. Uncropped SDS–PAGE images of pull-down assay showing interactions between MurJ and Lys$^{M}$ corresponding to Figure 2C, D**

(A) The original SDS–PAGE image of Fig. 2C. MurJ-His$_8$ and various Lys$^{M}$-GFP mutants were expressed and purified using a Ni-NTA column. The presence of MurJ-His$_8$ in the eluate was confirmed through CBB staining. Lys$^{M}$-GFP was detected via immunoblotting using an anti-His antibody. (B) The original SDS–PAGE image of Fig. 2D. Lys$^{M}$-His$_{10}$ and Lys$^{M}$-resistant MurJ mutants were expressed and purified using a Ni-NTA column. The presence of Lys$^{M}$-His$_{10}$ and MurJ-3×FLAG was confirmed via immunoblotting using anti-His and anti-FLAG antibodies, respectively.

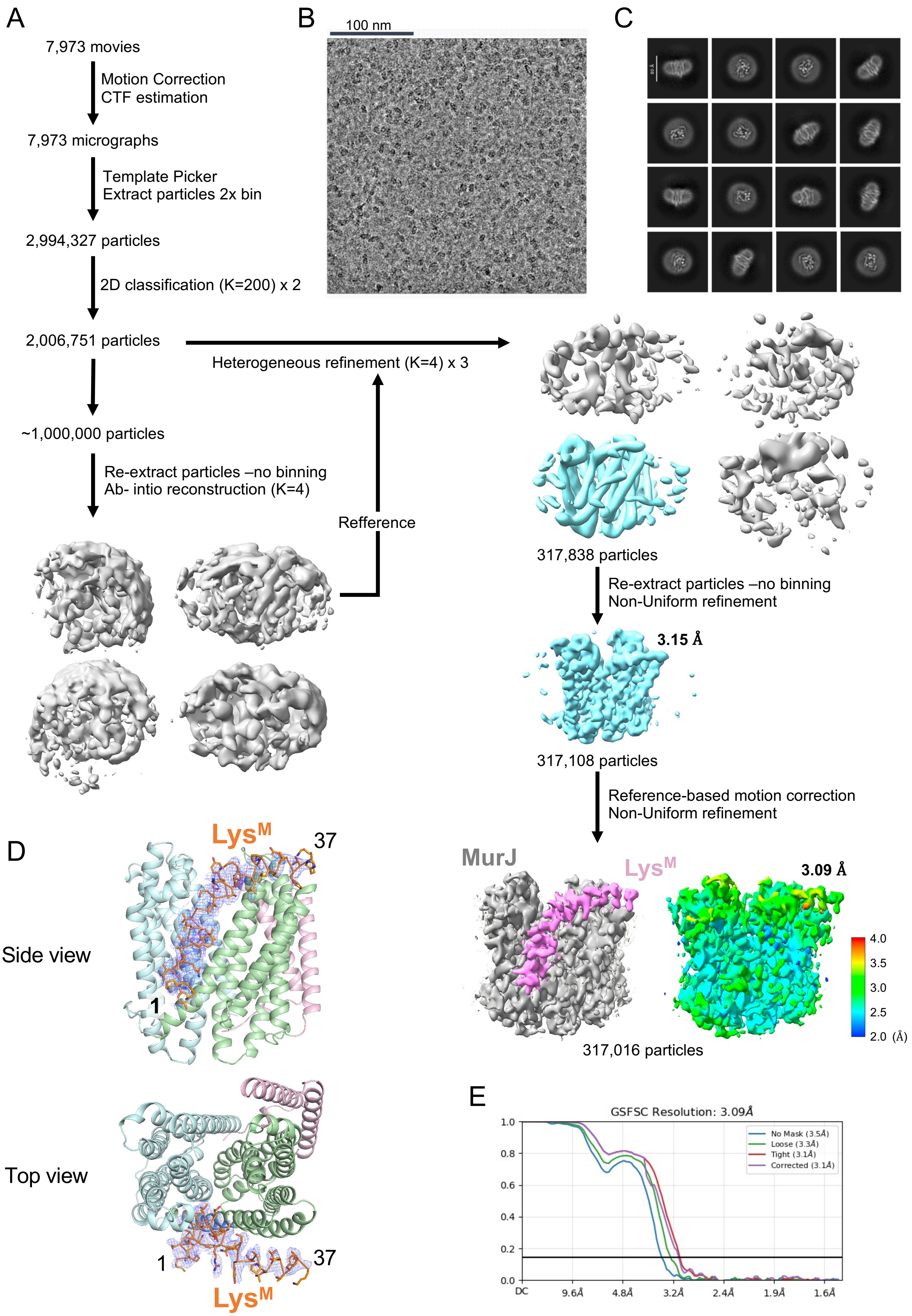


**Supplementary Figure 5. Cryo-EM data processing workflow for JM complex.**

(A) Image processing workflow for the JM complex. In the final reconstruction map, MurJ and $Lys^M$ are shown in gray and pink, respectively. Local resolution of the reconstructed map was estimated using CryoSPARC. (B) Representative electron micrograph of the nanodisc-reconstituted JM complex. (C) 2D class averages of the JM complex. (D) Cryo-EM map (displayed at 5σ) of $Lys^M$ in the JM complex and the JM complex model. (E) Gold-standard FSC curve used for global resolution estimation within CryoSPARC.

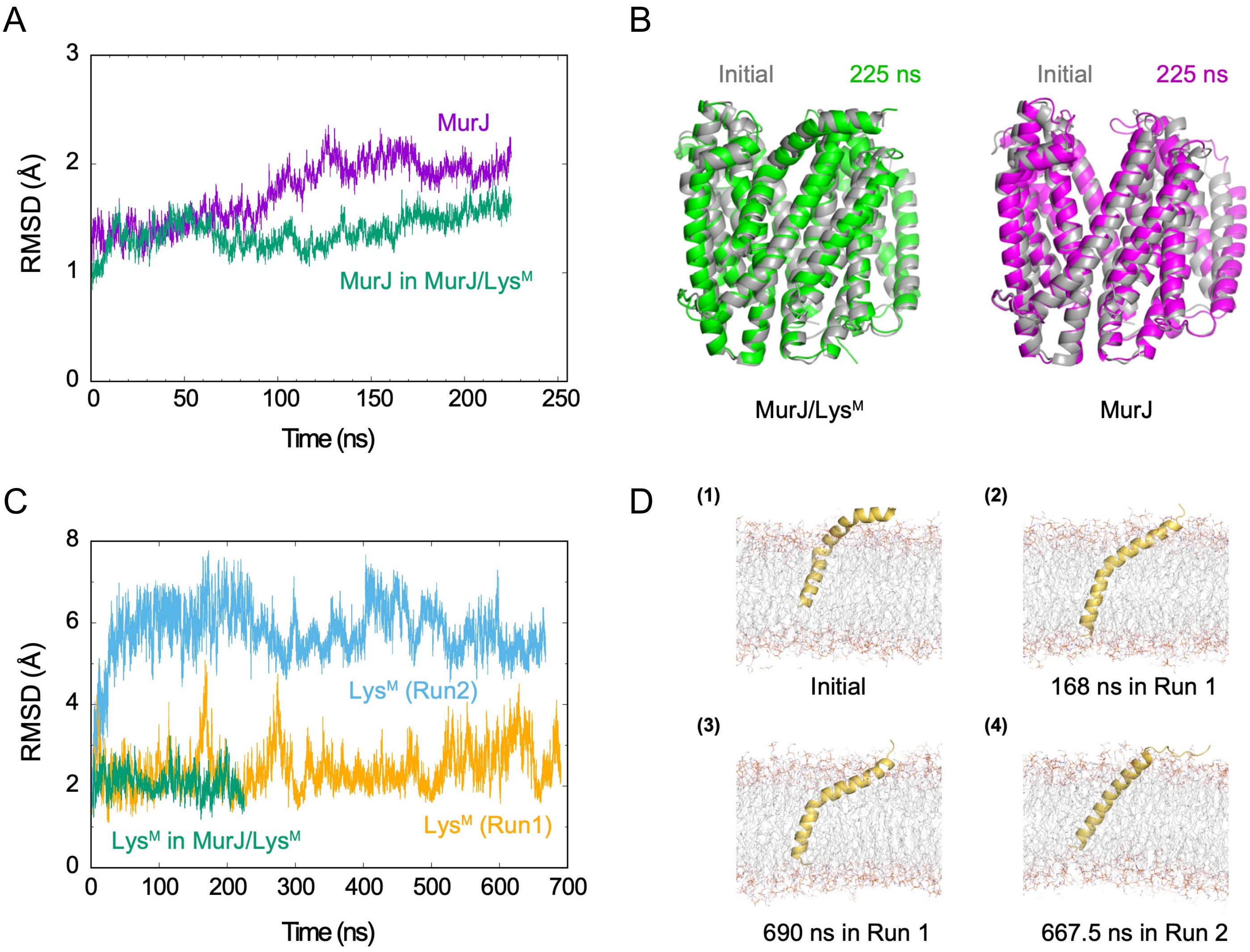


**Supplementary Figure 6. All-atom MD simulations of MurJ/Lys$^{M}$ complex, MurJ, and Lys$^{M}$ in membranes**
(A) Time evolution of the Cα-RMSD for MurJ in the MurJ/Lys$^{M}$ complex (green) and MurJ alone (magenta), relative to the initial structure. (B) Representative snapshots from the MD simulations of the MurJ/Lys$^{M}$ complex and MurJ alone, comparing the initial structure with the final conformation. (C) Time evolution of the Cα-RMSD for Lys$^{M}$ in the MurJ/Lys$^{M}$ complex (green) and Lys$^{M}$ alone (orange for Run1 and blue for Run2), relative to the initial structure. (D) Representative snapshots of the Lys$^{M}$ alone simulations. Initial structure (1) and conformations at 168 ns in Run 1 (2), 690 ns in Run 1 (3), and 667.5 ns in Run 2 (4).

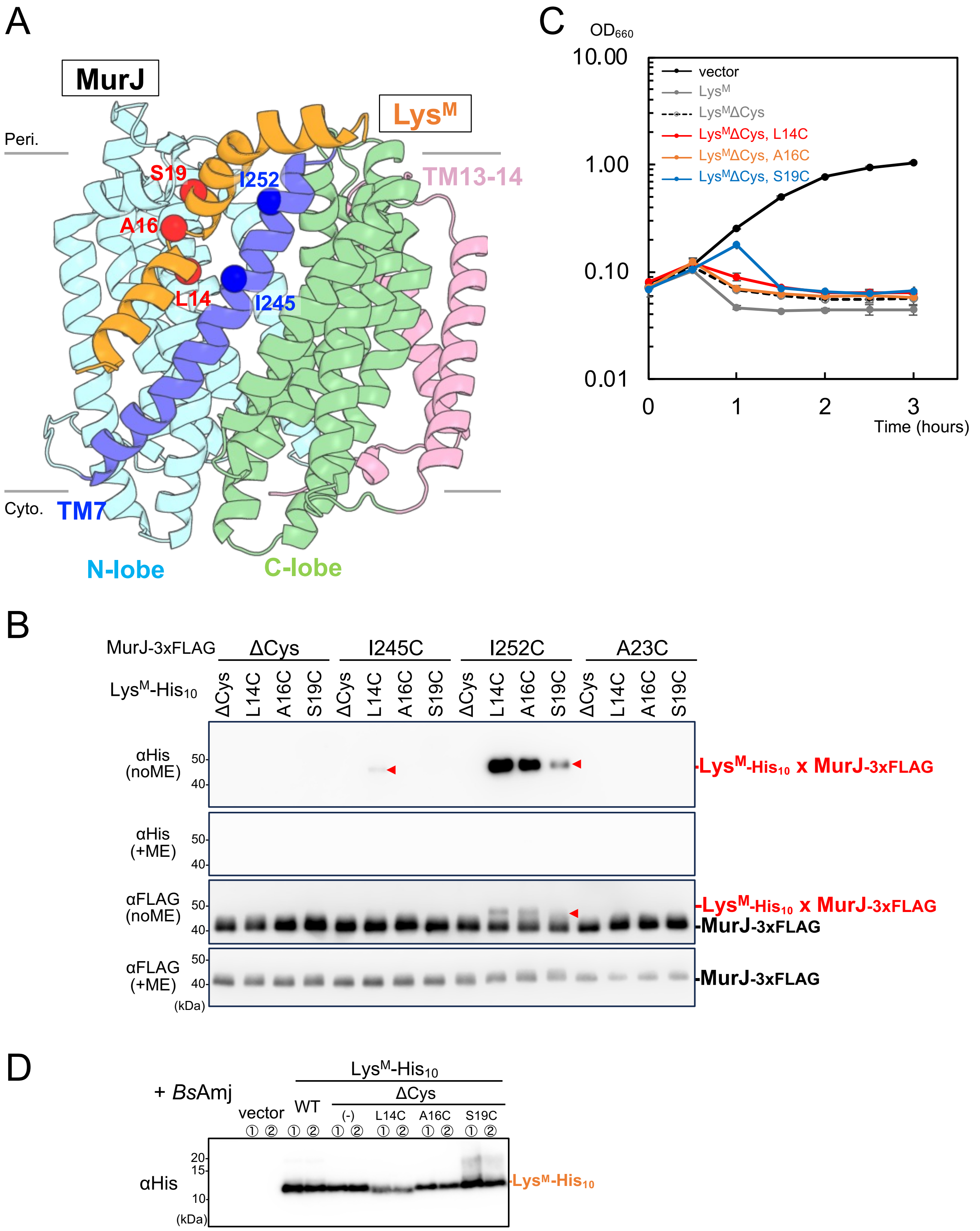


**Supplementary Figure 7.** ***In vivo*** **disulfide crosslinking between Lys$^{M}$ and MurJ.**

(A) Mapping of cysteine-substituted residues used for disulfide crosslinking onto the JM complex structure. The Cα atoms of Lys$^{M}$ residues (L14, A16, S19) and MurJ residues (I245, I252) are shown in red and blue, respectively. (B) *In vivo* disulfide bond formation between Lys$^{M}$ and MurJ. *E. coli* expressing Cys-substituted mutants of MurJ-3×FLAG and Lys$^{M}$-His$_{10}$ as indicated. After cultivation, samples were treated under oxidative conditions and analyzed using SDS–PAGE under reducing (+ME) or nonreducing (no ME) conditions, followed by immunoblotting with the indicated antibodies. Red arrowheads indicate crosslinked bands. (C) Lytic activity of Lys$^{M}$ Cys mutants. Growth of *E. coli* cells harboring pBAD18 (vector) or plasmids carrying the cloned *lysM* genes. Cells were grown in LB medium supplemented with 0.2% (w/v) L-arabinose. (D) Accumulation levels of Lys$^{M}$-His$_{10}$ Cys mutants in the presence of BsAmj. After 2-h induction with L-arabinose and IPTG, total cell lysates were acid-precipitated and analyzed using SDS-PAGE and immunoblotting with an anti-His antibody. This panel shows results from two samples tested under identical conditions.

**Table S1. Data collection and refinement statistics, related to Figure S5.**

| | *Ec*MurJ/Lys$^{M}$-ND (EMD-64256, PDB ID: 9UKV) |
|---|---|
| **Data collection and processing** | |
| Magnification | x60,000 |
| Voltage (kV) | 300 |
| Electron exposure (e−/Å$^2$) | 51.2 |
| Defocus range (μm) | -1.4 to -1.6 |
| Pixel size (Å) | 0.752 |
| Symmetry imposed | C1 |
| Initial particle images (no.) | 2,994,327 |
| Final particle images (no.) | 316,016 |
| Map resolution (Å) | 3.05 |
| FSC threshold | 0.143 |
| Map resolution range (Å) | 4.76 to 1.85 |
| **Refinement** | |
| Initial model used (PDB code) | Homology model and AlphaFold model |
| Model composition | |
| Non-hydrogen atoms | 4188 |
| Protein residues | 548 |
| Ligands | 0 |
| *B* factors (Å$^2$) | |
| Protein | 60.09 |
| Ligand | 0 |
| R.m.s. deviations | |
| Bond lengths (Å) | 0.003 |
| Bond angles (°) | 0.695 |
| **Validation** | |
| MolProbity score | 1.99 |
| Clashscore | 11.70 |
| Poor rotamer (%) | 2.95 |
| Ramachandran plot | |
| Favored (%) | 97.79 |
| Allowed (%) | 2.21 |
| Disallowed (%) | 0 |

**Table S2. Plasmids and Strains used in this study.**

| Plasmids | Vector | Encoded gene and description | Reference or source |
|---|---|---|---|
| pBAD33 | - | Expression vector, $P_{araBAD}$, $Cm^R$ | Guzman et al., 1995 |
| pKK568 | pBAD33 | *lys*$^M$ | This study |
| pKG74 | pBAD33 | *lys*$^M$-*pa* | This study |
| pKG75 | pBAD33 | *lys*$^M$(K2A)-*pa* | This study |
| pKG76 | pBAD33 | *lys*$^M$(Y3A)-*pa* | This study |
| pKG77 | pBAD33 | *lys*$^M$(I4A)-*pa* | This study |
| pKG78 | pBAD33 | *lys*$^M$(I5A)-*pa* | This study |
| pKG79 | pBAD33 | *lys*$^M$(N6A)-*pa* | This study |
| pKG80 | pBAD33 | *lys*$^M$(L7A)-*pa* | This study |
| pKG81 | pBAD33 | *lys*$^M$(F9A)-*pa* | This study |
| pKG82 | pBAD33 | *lys*$^M$(C10A)-*pa* | This study |
| pKG83 | pBAD33 | *lys*$^M$(V11A)-*pa* | This study |
| pKG84 | pBAD33 | *lys*$^M$(L12A)-*pa* | This study |
| pKG85 | pBAD33 | *lys*$^M$(L13A)-*pa* | This study |
| pKG86 | pBAD33 | *lys*$^M$(L14A)-*pa* | This study |
| pKG87 | pBAD33 | *lys*$^M$(V15A)-*pa* | This study |
| pKG88 | pBAD33 | *lys*$^M$(G17A)-*pa* | This study |
| pKG89 | pBAD33 | *lys*$^M$(D18A)-*pa* | This study |
| pKG90 | pBAD33 | *lys*$^M$(S19A)-*pa* | This study |
| pKG91 | pBAD33 | *lys*$^M$(I20A)-*pa* | This study |
| pKG92 | pBAD33 | *lys*$^M$(Y22A)-*pa* | This study |
| pKG93 | pBAD33 | *lys*$^M$(R23A)-*pa* | This study |
| pKG94 | pBAD33 | *lys*$^M$(V24A)-*pa* | This study |
| pKG95 | pBAD33 | *lys*$^M$(S25A)-*pa* | This study |
| pKG96 | pBAD33 | *lys*$^M$(Q26A)-*pa* | This study |
| pKG97 | pBAD33 | *lys*$^M$(Y27A)-*pa* | This study |
| pKG98 | pBAD33 | *lys*$^M$(L28A)-*pa* | This study |
| pKG99 | pBAD33 | *lys*$^M$(P30A)-*pa* | This study |
| pKG100 | pBAD33 | *lys*$^M$(L31A)-*pa* | This study |
| pKG101 | pBAD33 | *lys*$^M$(V32A)-*pa* | This study |
| pKG102 | pBAD33 | *lys*$^M$(D33A)-*pa* | This study |
| pKG103 | pBAD33 | *lys*$^M$(T34A)-*pa* | This study |
| pKG104 | pBAD33 | *lys*$^M$(F35A)-*pa* | This study |
| pKG105 | pBAD33 | *lys*$^M$(T36A)-*pa* | This study |

| | | | |
|---|---|---|---|
| pKG106 | pBAD33 | *lys*$^M$(K37A)-*pa* | This study |
| pKG126 | pBAD33 | *lys*$^M$-*gfp*-*his*$_6$ | This study |
| pKG164 | pBAD33 | *gfp*-*his*$_6$ | This study |
| pNL1 | pBAD33 | *lys*$^M$(N6A)-*gfp*-*his*$_6$ | This study |
| pNL4 | pBAD33 | *lys*$^M$(L13A)-*gfp*-*his*$_6$ | This study |
| pNL7 | pBAD33 | *lys*$^M$(L14A)-*gfp*-*his*$_6$ | This study |
| pNL10 | pBAD33 | *lys*$^M$(D18A)-*gfp*-*his*$_6$ | This study |
| pNL13 | pBAD33 | *lys*$^M$(I20A)-*gfp*-*his*$_6$ | This study |
| pNL15 | pBAD33 | *lys*$^M$(P30A)-*gfp*-*his*$_6$ | This study |
| pKG166 | pBAD33 | *lys*$^M$-*gfp* | This study |
| pKG211 | pBAD33 | *gfp* | This study |
| pNL35 | pBAD33 | *lys*$^M$(N6A)-*gfp* | This study |
| pNL39 | pBAD33 | *lys*$^M$(L13A)-*gfp* | This study |
| pNL23 | pBAD33 | *lys*$^M$(L14A)-*gfp* | This study |
| pNL28 | pBAD33 | *lys*$^M$(D18A)-*gfp* | This study |
| pNL29 | pBAD33 | *lys*$^M$(I20A)-*gfp* | This study |
| pNL32 | pBAD33 | *lys*$^M$(P30A)-*gfp* | This study |
| pBAD18 | - | Expression vector, $P_{araBAD}$, $Amp^R$ | Guzman et al., 1995 |
| pKG191 | pBAD18 | *lys*$^M$-*his*$_{10}$ | This study |
| pKG252 | pBAD18 | *lys*$^M$-(P30*amb*)-*his*$_{10}$ | This study |
| pKG253 | pBAD18 | *lys*$^M$-(L31*amb*)-*his*$_{10}$ | This study |
| pKG254 | pBAD18 | *lys*$^M$-(V32*amb*)-*his*$_{10}$ | This study |
| pKG255 | pBAD18 | *lys*$^M$-(D33*amb*)-*his*$_{10}$ | This study |
| pKG256 | pBAD18 | *lys*$^M$-(T34*amb*)-*his*$_{10}$ | This study |
| pKG257 | pBAD18 | *lys*$^M$-(F35*amb*)-*his*$_{10}$ | This study |
| pKG258 | pBAD18 | *lys*$^M$-(T36*amb*)-*his*$_{10}$ | This study |
| pKG259 | pBAD18 | *lys*$^M$-(K37*amb*)-*his*$_{10}$ | This study |
| pKG539 | pBAD18 | *lys*$^M$-(C10A)-*his*$_{10}$ | This study |
| pKG542 | pBAD18 | *lys*$^M$-(C10A, L14C)-*his*$_{10}$ | This study |
| pKG545 | pBAD18 | *lys*$^M$-(C10A, A16C)-*his*$_{10}$ | This study |
| pKG546 | pBAD18 | *lys*$^M$-(C10A, S19C)-*his*$_{10}$ | This study |
| pKG548 | pBAD18 | *lys*$^M$-(C10A, I20C)-*his*$_{10}$ | This study |
| pKG65 | pTSP1 | *EcmurJ*-*his*$_8$ | Kohga et al., 2022 |
| pSTV28 | - | Expression vector, $P_{lac}$, $Cm^R$ | Takara Bio |
| pKG537 | pSTV28 | *EcmurJ*-*3xflag* | This study |

| | | | |
|---|---|---|---|
| pNL71 | pSTV28 | *EcmurJ*(A59G)-*3xflag* | This study |
| pNL61 | pSTV28 | *EcmurJ*(F64L)-*3xflag* | This study |
| pNL73 | pSTV28 | *EcmurJ*(V229G)-*3xflag* | This study |
| pNL62 | pSTV28 | *EcmurJ*(M233L)-*3xflag* | This study |
| pNL68 | pSTV28 | *EcmurJ*(Q244P)-*3xflag* | This study |
| pNL65 | pSTV28 | *EcmurJ*(I248S)-*3xflag* | This study |
| pNL70 | pSTV28 | *EcmurJ*(I252F)-*3xflag* | This study |
| pKG554 | pSTV28 | *EcmurJ*(C314S, C419S)-*3xflag* | This study |
| pKG584 | pSTV28 | *EcmurJ*(C314S, C419S, A23C)-*3xflag* | This study |
| pKG593 | pSTV28 | *EcmurJ*(C314S, C419S, I245C)-*3xflag* | This study |
| pKG596 | pSTV28 | *EcmurJ*(C314S, C419S, I252C)-*3xflag* | This study |
| pTWV228 | - | Expression vector, $P_{lac}$, $Amp^R$ | Takara Bio |
| pRM1072 | pTWV228 | *Bacillus subtilis amj* | This study |
| pHM1550 | - | Expression vector, $P_{lac}$, $Spc^R$ | Miyazaki et al., 2022 |
| pRM1210 | pHM1550 | *Bacillus subtilis amj* | This study |

| Strains | Genotype | Reference or source |
|---|---|---|
| JM109 | *recA1*, *endA1*, *gyrA96*, *thi-1*, *hsdR17*(*rK- mK*+), *e14-* (*mcrA-*), *supE44*, *relA1*, Δ (*lac-proAB*)/F'〔*traD36*, *proAB*+, *lac Iq*, *lacZ*ΔM15〕 | Takara Bio |
| BL21(DE3) | *F-, ompT, hsdSB (rB-mB-), gal, dcm* | Takara Bio |
| BL21(DE3), ⊿recA | *F-, ompT, hsdSB（rB-mB-）, gal, dcm, ⊿recA, ⊿endA* | Nippon gene |
